# Drivers of diversification in individual life courses

**DOI:** 10.1101/103184

**Authors:** Raisa Hernández-Pacheco, Ulrich K. Steiner

## Abstract

Heterogeneity in life courses among individuals of a population influences the speed of adaptive evolutionary processes, but it is less clear how biotic and abiotic environmental fluctuations influence such heterogeneity. We investigate principal drivers of variability in sequence of stages during an individual’s life in a stage-structured population. We quantify heterogeneity by measuring population entropy, which computes the rate of diversification of individual life courses of a Markov chain. Using individual data of a primate population, we show that density regulates the stage composition of the population, but its entropy and the generating moments of heterogeneity are independent of density. This lack of influence of density on heterogeneity is neither due to low year-to-year variation in entropy nor due to differences in survival among stages, but due to differences in stage transitions. Our analysis thus shows that well-known classical ecological selective forces, such as density regulation, are not linked to potential selective forces governing heterogeneity through underlying stage dynamics. Despite evolution acting heavily on individual variability in fitness components, our understanding is poor whether observed heterogeneity is adaptive and how it evolves and is maintained. Our analysis illustrates how entropy represents a more integrated measure of diversity compared to the population structural composition, giving us new insights about the underlying drivers of individual heterogeneity within populations and potential evolutionary mechanisms.

## INTRODUCTION

Predicting ecological and evolutionary population dynamics requires an understanding of the diversity of life courses among individuals, since life courses shape demography, which in turn drives evolution (Caswell 2001, Metcalf and Pavard 2007). Much ecological and evolutionary analysis focuses on correlations among mean population-level factors (e.g., density-dependence) or expected life courses (e.g., optimization), but it is the variation among individual life courses that can, if adaptive, accelerate evolutionary change, or if neutral, can slow the pace of evolutionary change (Wright 1931, Crow and Kimura 1970, Steiner and Tuljapurkar 2012). To further our understanding of ecological and evolutionary processes within a population we need to examine the mechanisms underlying among-individual heterogeneity in life courses within- and among-environments (Lomnicki 1988, Bjørnstad and Hansen 1994, Novoseltsev et al. 2003, Lescroël et al. 2009).

Here, we aim at understanding how environmental variability changes individual level heterogeneity in life courses. We focus on population density as environmental factor and use population entropy to quantify diversification in individual life courses. Individual life courses can be formulated as a sequence of stages (stage trajectories) that starts with birth and ends with death (Caswell 2001). Trajectories progress through various developmental, reproductive, behavioral, morphological, physiological or similar stages. We can quantify the diversity of these life courses by computing a measure of entropy on a stage-structured population and describing the dynamics of stage transitions by a Markov chain (Tuljapurkar et al. 2009, Steiner et al. 2010). High rates of diversification in life courses, i.e. high entropy, have been found for numerous wild populations for which development and reproductive stages have been the focus (Tuljapurkar et al. 2009, Steiner et al. 2010, Plard et al. 2012); but these studies did not investigate the dependencies of entropy to either biotic or abiotic environmental fluctuations.

Our understanding remains limited of how heterogeneity among individual life courses is influenced by changes in survival and reproductive stage dynamics. In particular, we lack understanding on how changes in environmental and ecological factors such as population density influence heterogeneity among individual life courses. Changes in density may influence individual life courses in contrasting ways. Either, as the population grows, the increased density drives individuals to a certain “optimal” life course, thereby reducing entropy. Alternatively, increased density might lead to strong intraspecific competition within stages forcing individuals to explore new niches (Rubenstein 1981) by diversifying life courses. As a result of the second expectation entropy will increase. The density-dependent mechanism underlying the unification or diversification of life courses (variation in entropy) may depend on current annual density (Boyce 1984), past annual density (e.g., time-lag effects) (Magnuson 1990), or density at birth (e.g., cohort effects) (Beckerman et al. 2002, Nussey et al. 2007). In this way, we may expect across-year differences (period effects) in biotic factors (e.g., density) (Tuljapurkar et al. 2009) or even cohort selection (Vaupel and Yashin 1985) (e.g., density at birth) to influence entropy; the former factor by increasing variability in survival and the latter by decreasing it. Which of these alternative hypotheses is supported might depend on the stage type assessed, e.g., development, reproduction, behavior, morphology, physiology, or geographic location.

In this study, we evaluate the effects of population density on the variability in individual life courses of Cayo Santiago rhesus macaques by computing population entropy for annual stage-structured population matrix models and using it as a proxy of annual diversification in individual life courses. Here, stages are defined as development and reproductive stages. We also assessed whether changes in survival, reproduction, age, and stage dynamics influenced population entropy. Finally, we identify which stage transition contributed the most to changes in entropy by performing perturbation analyses on stage-transitions. As the properties of a dynamic life history depend on all the transition rates, our assessment based on entropy presents a more integrated measure of diversity, and thus gives us new insights about heterogeneity within populations (Tuljapurkar et al. 2009). Our analysis reveals that changes in density influence the observed stage distribution in the population; however, the diversification in individual life courses (entropy) is independent of the mechanisms underlying structural composition (stage distributions).

## METHODS

Based on a free-ranging rhesus monkey population, we parameterized for each period (year) a stage-structured matrix model. For those resulting 40 matrixes we estimated the population entropy and explored its relationship to population density and other additional variables such as survival, age, and birth cohort.

### The Cayo Santiago rhesus macaque population

Cayo Santiago is a 15.2-ha island located 1 km off the south-eastern coast of Punta Santiago, Puerto Rico (18° 09’ N, 65° 44’ W) that has served as habitat for free-ranging rhesus monkeys (*Macaca mulatta*) since 1938 (Carpenter 1972, Rawlins and Kessler 1986). All individuals are descendent of the 409 Indian-origin founders. Since then, the colony has been maintained under semi-natural conditions for behavioral and non-invasive research (Kessler and Rawlins 2016). Monkeys forage on vegetation and are provisioned with commercial monkey chow. The daily amount of food provisioned is estimated per capita as 0.23 kg/animal/day, an amount that is chosen to avoid food limitation. Food is distributed among three open feeding stations located at different sites of the island. Rainwater is collected in catchments on the island, stored in concrete or fiberglass cisterns, and chlorinated prior to distribution at automatic watering stations. Monkeys have free access to both feeding sites and watering. Veterinary intervention is minimal except during the annual trapping period when yearlings are captured, marked for identification, and physical samples are collected. Each monkey has a unique identification tattoo on its chest and thigh. Information on each monkey includes identity, date of birth, sex, maternity, maternal lineage, social group, and date of death. Due to management purposes, a portion of the population could be permanently removed from the island during a particular trapping season. Demographic data on removed individuals is equally recorded with a date of removal instead of date of death. A monthly census is generated from daily observations made of the entire population by the Caribbean Primate Research Center (CPRC) staff. This population is ideal to assess the drivers of heterogeneity in individual life courses, as detailed data of every individual in the population is available.

Cayo Santiago monkeys mate and reproduce once a year in a synchronized fashion (>72% of births within a 3-month period) demarcating an annual birth season (Hernández-Pacheco et al. 2016a). Females start reproducing at 3 years of age with 10-20% of the 3-year old and > 30% of 4-year-olds and older females giving birth (Hernández-Pacheco et al. 2013). No birth has been recorded after females reach 23 years of age and the oldest recorded female died at 31 years of age (Hernández-Pacheco et al. 2013). Changes in density are clearly identifiable because it is a closed island population. More importantly, this primate population exhibits a high annual population growth rate (*λ* > 1.10) (Rawlins and Kessler 1986, Blomquist et al. 2011, Kessler et al. 2015), and exists frequently at relatively high densities where population dynamics are regulated by reduced reproduction (Hernández-Pacheco et al. 2013, Hernández-Pacheco et al. 2016b).

### Demographic model parameterization

Our analysis is based on the 40-year period from 1973 to 2013. Annual censuses are based on daily census data of 3,901 females. We used a birth-pulse model employing post-breeding censuses including female data only and discrete time stage-structured matrix models with annual intervals (periods) (Caswell 2001). Even though the timing of births has shifted in Cayo Santiago rhesus macaques (Hernández-Pacheco et al. 2016a), we kept the annual structure in our analysis, which did not lead to overlapping breeding periods. We defined stages as development and reproductive stages. In a given reproductive season, we classified <3 year-old females as sexually immature individuals (stage I). Three-year-old and older females were classified into one of three following reproductive stages: (1) non-breeders (stage NB), females that did not give birth in a given reproductive season; (2) failed breeders (stage FB), females that gave birth but their offspring did not survive to 1 year of age (independence); and (3) successful breeders (stage B), females that gave birth and their offspring survived to >1 year of age. We right censored post-reproductive females (>23 year olds; N = 75) in order to avoid an overestimation of non-breeders.

Using these data, we parameterized 40 matrix models, one for each annual period *t*, with stage-transition matrix, **P**_*t*_, and stage-specific survival rates,

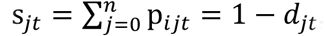
where *d*_*jt*_ is stage-specific mortality in stage *j* and period *t*. After the first 3 years of life spent in stage I, surviving females transitioned to and thereafter among one of the three reproductive stages (NB, FB, or B) until death, until reaching post-reproductive stage, or until being removed (culled) from the island. For each matrix model, we estimated the asymptotic population growth rate (*λ*_1_), the stable age distribution (***w***), and the reproductive value vector (***v***) from the dominant eigenvalue of **P**_*t*_ and its corresponding right and left eigenvectors, respectively (Caswell 2001). We carried out all analyses using R.3.0.1, (R Development Core Team). In order to determine the variables influencing entropy, as well as whether vital rates are density-dependent, we fitted linear and quadratic models and carried out model selection using Akaike’s Information Criterion (AIC) (Burnham and Anderson 2002). We interpreted a difference of two or more between AIC values to indicate that two models differed in their support (Burnham and Anderson 2002). When comparing models with different response variables, we present the residual deviance. To justify our model selection, we evaluated the overall fit of our basic models by generating a single fully time dependent stage-structured model and estimated the goodness of fit (GOF) of this model using the bootstrapping function of program MARK (White et al. 2001). Even though such a single model deviates slightly from the 40 models in its (stationary) stage dynamics, we based our judgment of goodness of fit on this model, which revealed little overdispersion (ĉ=1.057).

### Patterns in reproductive stage distribution

We know that the proportion of births with respect to the total number of adult females (mean fertility) depends on density in this population (Hernández-Pacheco et al. 2013). In order to examine whether our annual stage structured models accurately represent the density-dependent dynamics in stage structure, we compared the observed stage distributions to the stationary stage distributions estimated by our annual models (another measure of goodness-of-fit). In order to avoid underestimation of a particular annual stage distribution in a period with adult female removal (total of 11 periods), the annual proportion of females was estimated using the total number of females alive at period *t* after censoring removed adult females during the same period. Although selective culling in the population has been practiced, adult females have never been the main target or their reproductive stage (Hernández-Pacheco et al. 2016b). We determined mean annual fertility (*F*_*t*_) as the proportion of successful breeders (B) relative to the total number of adult females (NB, FB, B) during the birth season of period *t*. In our annual models, only stage B contributes to reproduction, while stage FB contributes to an absorbing stage (dead infants). We assessed whether adult density explained the observed proportion of B, NB, and FB females during the same period (density-dependence on reproductive stages). Given the fact that social interactions are thought to be the main mechanism underlying density-dependence in this population (Hernández-Pacheco et al. 2013), we also included the total number of adult males (same age range as females was used), when assessing density-dependence in female reproduction. We repeated this density-dependent analysis for each reproductive stage-specific survival rate (s_*jt*_) and transition probabilities among reproductive stages (p_*ij*_). Density dependence might not always be showing an immediate effect, hence we investigated if effects might be delayed by evaluating density-dependence using the number of adult individuals at period *t*-1 with the proportion of B, NB, and FB at period *t* (time-lag effects). Time-lag effects were also evaluated in s_*jt*_.

### Variability in life courses: population entropy

An individual life course can be viewed as a sequence of reproductive stages terminating with death. Population entropy *H* describes the rate at which the diversity of these individual stage trajectories (individual life courses) increases with age (Table I, II) (Pielou 1977, Tuljapurkar et al. 2009). We quantified this variation using the absorbing matrix **R** given the transition probabilities in matrix **P** weighted by its corresponding quasi-stationary proportions π (Table I, II) (Matthews 1970, Steiner et al. 2010; Online Appendix A). Entropy *H* increases as the probability of individuals becomes more even to transition between any of the reproductive stages. Entropy is limited between 0, full deterministic stage sequences, and its maximum value ln *K*; where *K* equals the number of reproductive stages (Tuljapurkar et al. 2009). We scaled entropy to its maximum value as to *H*/*H*_*max*_; hence, scaled *H* is bounded between, 0 and 1. Here, we extend the current use of *H* to annual matrices by using it as a proxy of the annual rate of diversification among individuals. From each **P**_*t*_, we parameterized annual absorbing matrices, **R**_*t*_ and estimated its corresponding entropy, *H*_**R***_t_*_ The latter allows assessing the influence of survival on heterogeneity.

**Table I.** Parameter index

| Parameter |  |
| --- | --- |
| $t$ | Period (year) |
| $F_t$ | Mean fertility in year $t$ |
| $i, j$ | Stage |
| $s_i$ | Stage-specific survival rate |
| $d_j$ | Stage-specific mortality |
| $\mathbf{A}$ | Mean 4x4 projection matrix |
| $\mathbf{P}$ | Submatrix of $\mathbf{A}$ based on adult stages only (mean 3x3 stage-transition matrix across periods $t$ with stage-specific survival rates, $s_j$, $\sum_{i=0}^K p_{ij} = 1 - s_j$) |
| $p_{ij}$ | Transition probabilities of $\mathbf{P}$ |
| $\mathbf{P}_t$ | Stage-transition matrix for year $t$ with stage-specific survival rates, $s_{jt}$ |
| $\mathbf{w}$ | Stable stage distribution of $\mathbf{P}$ (scaled right Eigen vector corresponding to dominant eigenvalue, as to $\sum_{i=0}^K w_i = 1$) |
| $\mathbf{v}$ | Reproductive value vector of $\mathbf{P}$ (scaled left Eigen vector corresponding to dominant eigenvalue, as to $v_i = 1$) |
| $\mathbf{P}'$ | Non-absorbing stage-transition matrix with $\sum_{i=0}^K p_{ij} = 1$, i.e. excluding survival and mortality |
| $\mathbf{W}$ | Diagonal matrix with elements corresponding to $\mathbf{w}$ (see Table II) |
| $\mathbf{R}$ | Absorbing 3x3 stage-transition matrix based on $\mathbf{P}$ (see Table II) |
| $\mathbf{R}_t$ | Period-based absorbing stage-transition matrix |
| $\lambda_1$ | Dominant Eigen value of $\mathbf{P}$ |
| $H$ | Population entropy |
| $H_{\mathbf{R}_t}$ | Entropy of matrix $\mathbf{R}_t$ |
| $H_{\mathbf{P}'_t}$ | Entropy of matrix $\mathbf{P}'_t$ (excluding survival) |
| $\mathbf{R}_c$ | Cohort-based $\mathbf{R}$ |
| $H_{\mathbf{R}_c}$ | Entropy of $\mathbf{R}_c$ |
| $K$ | Row or column number of matrix $\mathbf{R}$ |

**Table II:** Measuring population entropy in an absorbing Markov chain

| Quantity | Definition |
| --- | --- |
| Dominant Eigen value of $\mathbf{P}$ | $\lambda_1$ |
| Matrix $\mathbf{R}$ | $\mathbf{R} = \frac{1}{\lambda_1} \mathbf{W}^{-1} \mathbf{P} \mathbf{W}$ |
| Elements of matrix $\mathbf{R}$ | $\mathbf{R}_{ij} = \frac{1}{\lambda_1} \frac{\mathbf{P}_{ij} \mathbf{W}_j}{\mathbf{W}_i}$ |
| Quasi Stationary distribution | $\pi_i^T = \pi_i^T \mathbf{R} = \mathbf{v}^T \mathbf{W}$ |
| Entropy in absorbing Markov chain | $H = - \sum_{i=1}^K \sum_{j=1}^K \pi_i \mathbf{R}_{ij} \log \mathbf{R}_{ij}$ |
| Entropy in nonabsorbing Markov chain | $H = - \sum_{i=1}^K \sum_{j=1}^K w_i \mathbf{P}_{ij}^T \log \mathbf{P}_{ij}^T$ |

Superscript T refers to transpose

### Heterogeneity generated by density

In this study, we aim at understanding the mechanisms underlying the structural changes that generate the variability in individual life courses. First, we tested for density effects on entropy by evaluating whether the total number of adults in a given period *t* influenced *H*_**R***_t_*_. As individuals experience different densities among years, changes in *H* driven by a change in density would be captured in our Markovian models (annual matrices). In order to support further this assumption, we evaluated across year (period) trends in density as well as lag-time effects, and determined whether *H*_**R***_t_*_ was correlated with time (periods effects). We also tested whether our models paralleled the observed data by regressing the annual proportion of females in the different reproductive stages with the predicted proportion of the stable stage distribution (SSD), ***w*** (Table I), given by each annual model.

Second, we evaluated whether the annual proportion of adult females, which is density-dependent (see Results), influenced *H*_**R***_t_*_ Similarly, we assessed the influence of the variability in transition probabilities (which might also depend on density) on *H*_**R***_t_*_. Finally, we assessed whether population density influences variability in life courses through birth cohort effects (density at birth) rather than across cohorts for each period. For this, we divided the female population into (birth) cohorts. For each cohort, we tracked the life course of all members and parameterized with these data one matrix ***R***_***c****_t_*_ and its corresponding entropy, *H*_**R**c_ (Table I). We tested whether ***R***_***c****_t_*_ correlated with adult density at period *t*. We also tested the cohort-based *H*_Rc_ against the corresponding entropy of the period-based transition matrix, *H*_**R***_t_*_ (e.g.,

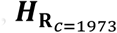
 versus

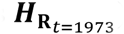
). This analysis was carried out until 1999 in order to maximize the number of ages in each cohort and prevent bias due to right censoring.

### Heterogeneity generated by age

Heterogeneity and diversification might also be driven by age, i.e. senescence in reproduction and survival. Such senescence has been described for this population (Hoffman et al. 2010) and we also reveal it when we analyzed a stage-structured model across time (time independent stage-age structured model). Because of sparse data and convergence problems as well as estimability issues we could not fit fully age-time-stage-structured models. We reran part of our analysis by truncating all ages 17 and older and revealed that the results are robust to this truncated data (Online Appendix B). The age 17 was chosen as it is the onset of senescence in survival and reproduction. We conclude that heterogeneity and stage dynamics and our findings are not particularly driven by old individuals. Thus, we present results on all reproductive females (3-23 years of age). For conciseness we do not present any detailed results on the age specific analyses.

### Heterogeneity generated by survival

The variation in *H*_**R***_t_*_ could also be driven by survival differences among stages rather than by stage transitions alone. We evaluated the effects of survival differences on entropy by comparing *H*_**R***_t_*_(absorbing Markov chain based, with death as the absorbing stage) to the entropy of the corresponding transition matrix **P**′_***t***_ that does not account for stage survival (*H*_**P**′*t*_) (non-absorbing Markov chains) (Tuljapurkar et al. 2009, Steiner et al. 2010) (Table I). In this way, any difference between both annual estimates of *H* (*H*_**R***_t_*_ and *H*_**P**ʹ*t*_) reveals the influence of survival differences among stages on the variability in life courses.

### Sensitivity analysis of entropy

To illustrate the sensitivity of *H* to a change in the stage-specific transition probabilities (p_*ij*_), we perturbed p_*ij*_ in mean matrix **P** allowing survival to remain constant, and computed *H* for each perturbed matrix. Mean matrix **P** was defined as the mean stage-transition matrix across all periods (Table I). Specifically, we modeled three different scenarios and tested each simulated outcome with the *H*_**R***_t_*_ from the observed data. For this, we increased the probability of stasis in NB (p_NB,NB_) from 0 to *a*, where *a* equals the sum of the other transition probabilities being perturbed. As NB stasis increased, a proportional decrease in i) the transition from NB to FB (i.e. more FB move to and stay as NB); ii) the transition probability from NB to B (i.e. more B move to and stay as NB); and iii) both the transition probabilities from NB to FB and NB to B simultaneously (i.e. more B and FB move to and stay as NB) was performed. We present simulations on NB transitions, exclusively, as NB holds the highest proportion in the stable stage distribution (***w***) and thus, holds a higher potential for influencing *H* (see Results).

## RESULTS

### Density-dependence in reproduction

We found that the annual distribution of reproductive stages was influenced by population density, which supports previous results describing density-dependent dynamics in this population (Table IV; Figure 1) (Hernández-Pacheco et al. 2013). For our monkey population, the proportion of B, and thus *F*_*t*_, peaked at intermediate densities of adult individuals (~450 individuals) (Figure 1, open circles), while NB and FB showed opposite patterns (Figure 1, closed circles and crosses, respectively). Hence, annual changes in density influence the annual population structure of the population by suppressing reproduction or increasing unviable births at both low and high densities. Transition probabilities were independent of population density (Figure 2, Table AC I). Similarly, survival rates were independent of changes in density (Table AC II, Figure AD 1).

**Table III.** Mean projection matrix **A** and submatrix **P** of female macaques across all years

| **Period $t+1$** | **Reproductive stages at period $t$** |  |  |  |
| --- | --- | --- | --- | --- |
|  | **I** | **NB** | **FB** | **B** |
| **I** | 0.746 | 0 | 0 | 0.272 |
| **NB** | 0.214 | *0.605* | *0.498* | *0.628* |
| **FB** | 0.001 | *0.041* | *0.076* | *0.039* |
| **B** | 0.002 | *0.297* | *0.268* | *0.304* |

Note: I = Immature, NB = Non-breeders, FB = Failed Breeders, B = successful Breeders. Transition probabilities in italics represent the 3x3 submatrix **P** (grey section) used to estimate matrix **R**. Matrix columns do not sum up to 1 as stage-specific mortality is $> 0$ .

**Table IV:** Influence of population density on the observed proportion of adult female macaques

| **Reproductive stage** | **Slope** | **Quadratic term** | **df** | **AIC** |
| --- | --- | --- | --- | --- |
| NB null model (intercept only) | - | - | 2 | -137.11 |
| NB (linear model) | 0.0001 | - | 3 | -141.14 |
| NB (curvilinear) | -0.0003 | 0.0000003 | 4 | **-144.19** |
| NB (linear w/ time-lag model) | 0.00007 | - | 3 | -134.47 |
| B Null model | - | - | 2 | -139.62 |
| B (linear model) | -0.00006 | - | 3 | -140.34 |
| B (curvilinear) | 0.0005 | -0.0000005 | 4 | **-149.73** |
| B (linear w/ time-lag model) | -0.00006 | - | 3 | -137.82 |
| FB null model | - | - | 2 | -212.10 |
| FB (linear model) | -0.00003 | - | 3 | -214.24 |
| FB (curvilinear) | -0.0002 | 0.0000002 | 4 | **-218.71** |
| FB (linear w/ time-lag model) | -0.00002 | - | 3 | -209.90 |

**Figure 1:**
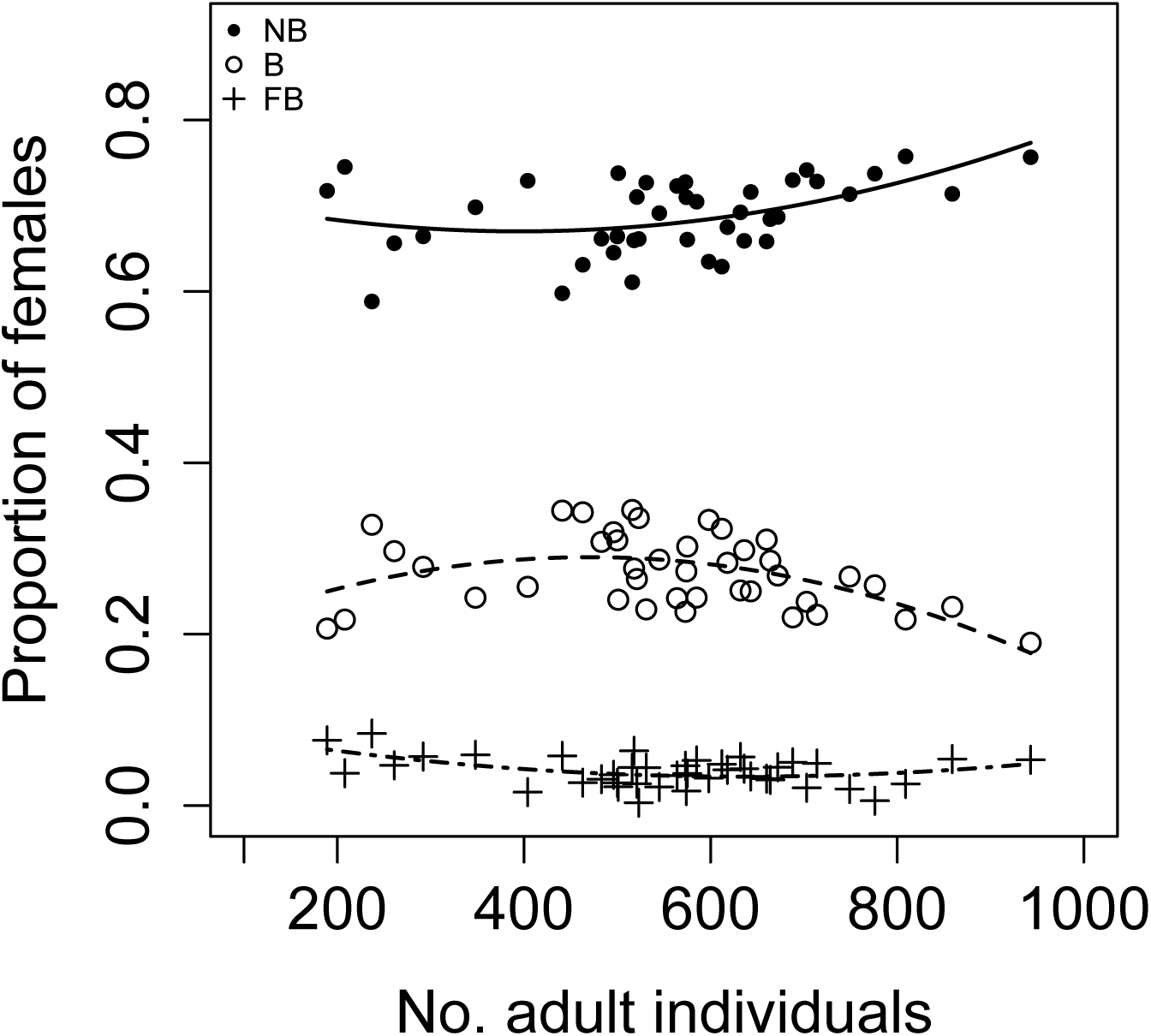
Density-dependent relationship between the annual proportion of nonbreeder (NB; close circles), breeder (B; open circles), and failed breeder (crosses) females and the total number of adult macaques in the population. Note that the annual proportion of breeders equals to the annual fertility rate (number of successful babies/number of adult females).

**Figure 2:**
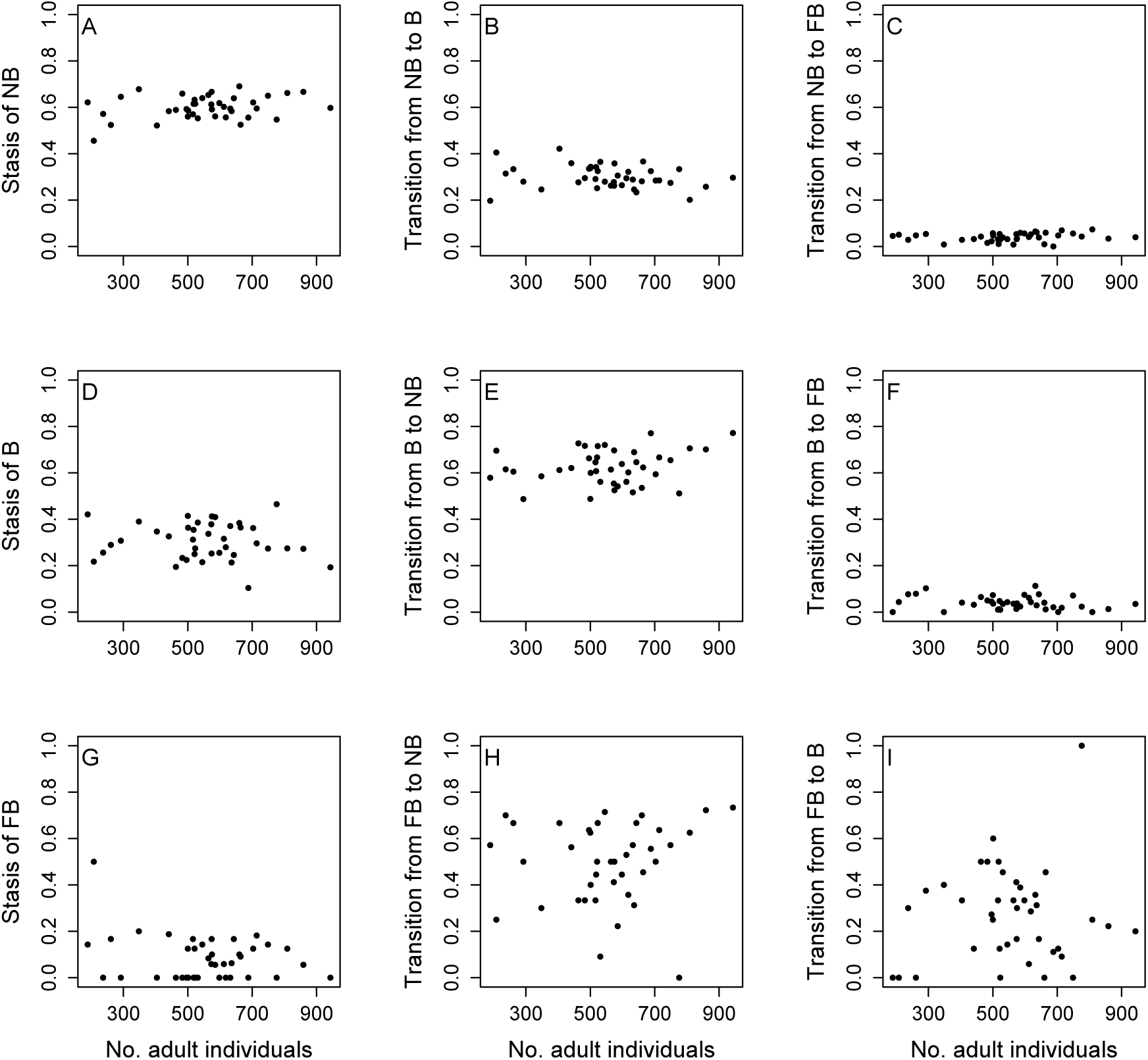
Transition probabilities as a function of the number of adult macaques.

### Patterns in reproductive stage dynamics

Our mean stage-structured model describes both survival and reproductive-stage dynamics (4x4 projection matrix **A**, Table III). For any given reproductive stage individuals have nonzero transition probabilities, indicating that two immature individuals that make it to age three will likely have distinct reproductive trajectories beyond that age. The scaled entropy we compute for the 3x3 mean submatrix **P** across all years (Table III, gray area) equals *H* = 0.71. If we estimate entropy for each period, the annual entropy *H*_**R**__*t*_ varied between 0.54 and 0.80, but there was no trend across periods (1973 to 2013) (Figure 3A). Furthermore, the stable stage distribution (SSD) (***w***) computed from **P**_*t*_ (Figure 3B) correlated with the observed stage-specific distribution of mature females (Figure 3D) (R^2^ = 0.82, 0.95, 0.79 for NB, FB, B, respectively), indicating that our annual models that are based on stable stage theories accurately represent the observed changes in stage distributions (Online Appendix E). Hence, we do not require to call transient dynamic models for which we would not know how to formally assess individual heterogeneity. The mean stable stage distribution **
*w*
** we estimate is biased towards NB females with 23.2% I, 51.0% NB, 3.18% FB, and 22.7% B.

**Figure 3:**
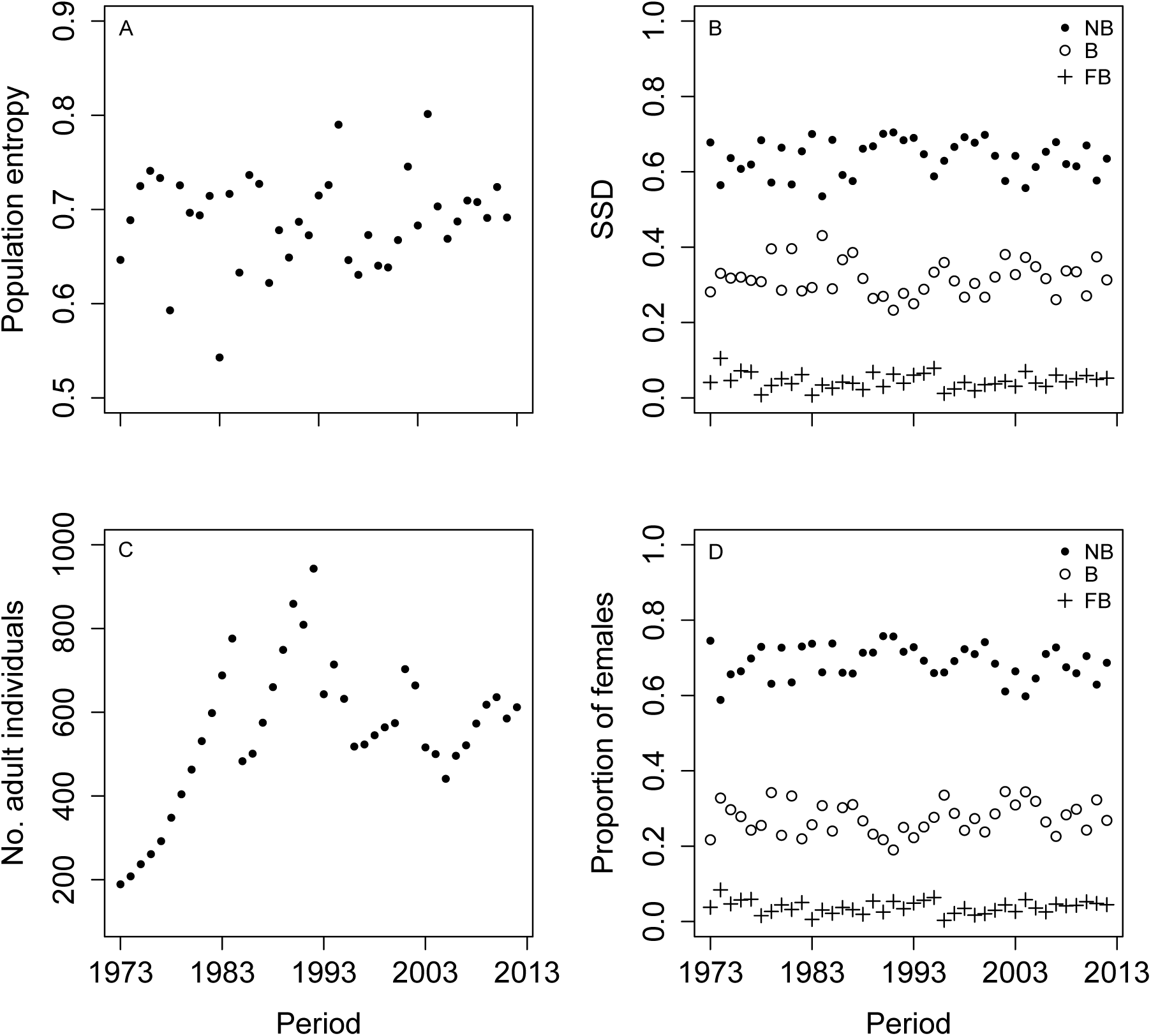
Variability in (A) population entropy, (B) stable stage distribution (SSD), (C) annual number of adults and (D) observed proportion of adult female rhesus macaques from 1973-2013.

### Density-dependence, survival, and population entropy

Our findings show that the number of adult individuals varied from 189 to 943 (Figure 3C) (females varied from 92 to 460 and males varied from 97 to 483) but surprisingly density did not influence *H*_**R**_*t*__ (Table V, Figure 4). Similarly, density at birth did not influence the observed changes in *H*_**R***_c_*_ (Table V and Table VI), and we did not find a correlation between *H*_**R***_t_*_ and *H*_**R***_c_*_ (cohort effects) (Online Appendix F). Only transition probabilities of NB and B influenced *H*_**R***_t_*_ (Figure 5).

**Figure 4:**
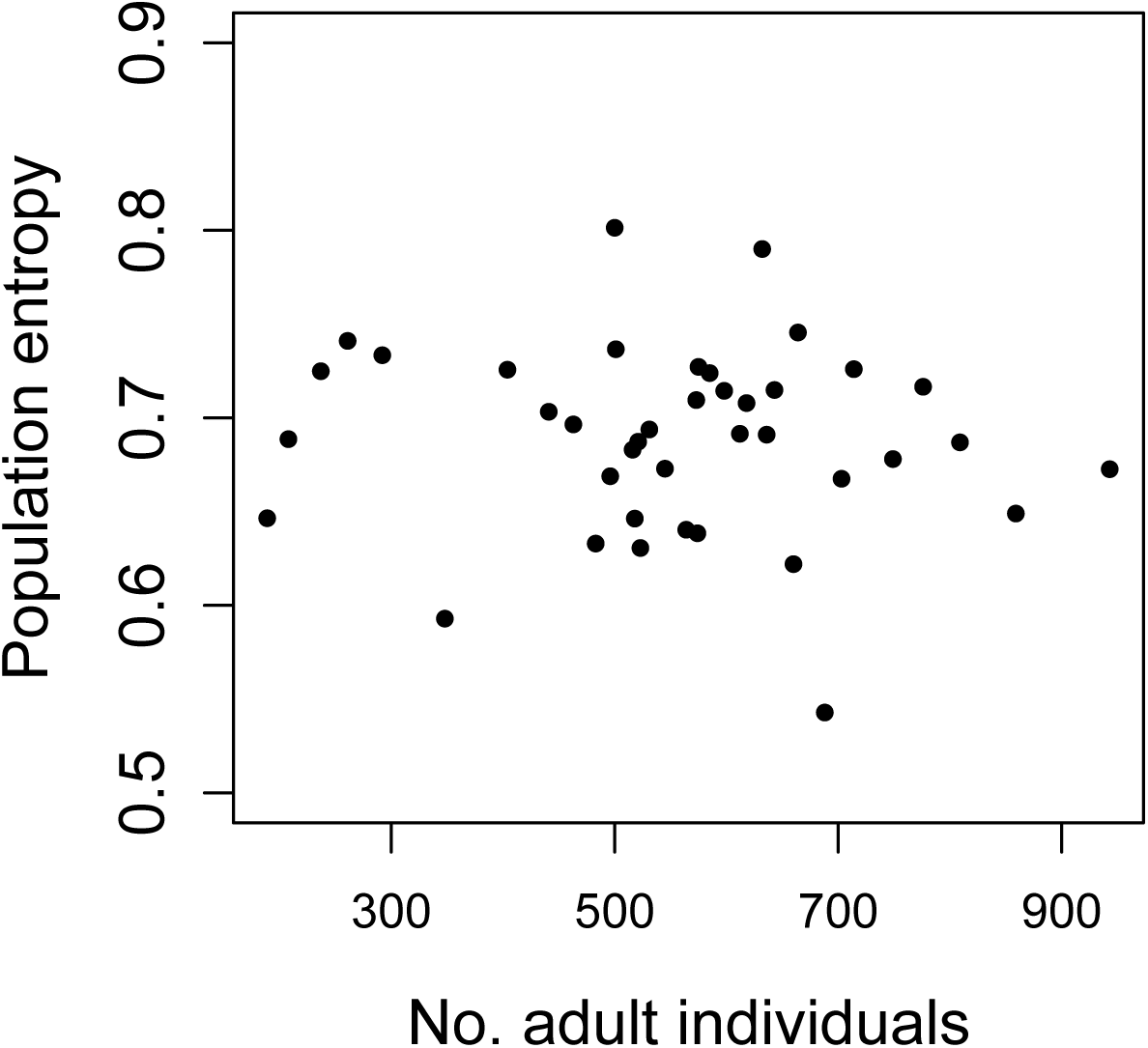
Annual population entropy *H*_**R***_t_*_ as a function of the number of adult macaques.

**Table V:** Variability in *H*_**R***_t_*_ as a function of changes in female proportion, survival, reproduction, transition probabilities, and total adult density.

| **Factors** | **Slope** | **Quadratic term** | **df** | **AIC** |
| --- | --- | --- | --- | --- |
| Null model (intercept only) | - | - | 2 | -123.72 |
| Proportion FB (curvilinear) | 5.228 | -39.820 | 4 | **-155.45** |
| Transition of NB to FB (curvilinear) | 4.352 | -32.715 | 4 | **-152.66** |
| Proportion NB (linear) | -0.673 | - | 3 | **-139.44** |
| Transition of B to FB (linear) | 0.918 | - | 3 | **-134.51** |
| Transition of B to NB (linear) | -0.340 | - | 3 | **-133.99** |
| NB stasis (curvilinear) | 4.077 | -3.826 | 4 | **-131.71** |
| B stasis (curvilinear) | 1.294 | -1.774 | 4 | **-128.94** |
| Proportion B (linear) | 0.419 | - | 3 | **-127.02** |
| FB survival (linear) | 0.123 | - | 3 | **-125.97** |
| NB survival (curvilinear) | 23.840 | -12.657 | 4 | -123.63 |
| Transition of NB to B (linear) | 0.274 | - | 3 | -124.78 |
| Transition of FB to B (linear) | 0.059 | - | 3 | -124.10 |
| B survival (linear) | 0.317 | - | 3 | -122.94 |
| Density (linear) (see Fig. 4) | -0.00003 | - | 3 | -122.05 |
| Time-lag in density (linear) | -0.00002 | - | 3 | -121.87 |
| FB stasis (linear) | -0.017 | - | 3 | -121.77 |
| Transition of FB to NB (linear) | -0.008 | - | 3 | -121.75 |

**Table VI:** Cohort-based analysis on *H*

| **Model** | **Slope** | **DF** | **Deviance** | **AIC** |
| --- | --- | --- | --- | --- |
| $H_{Rt} \sim 1$ | - | 26 | 0.0747 | **-78.42** |
| $H_{Rt} \sim \text{Density}$ | -0.00003 | 25 | 0.0739 | -76.71 |
| $H_{Rc} \sim 1$ | - | 26 | 0.0387 | **-96.19** |
| $H_{Rc} \sim \text{Density}$ | 0.00002 | 25 | 0.0381 | -94.57 |

**Figure 5:**
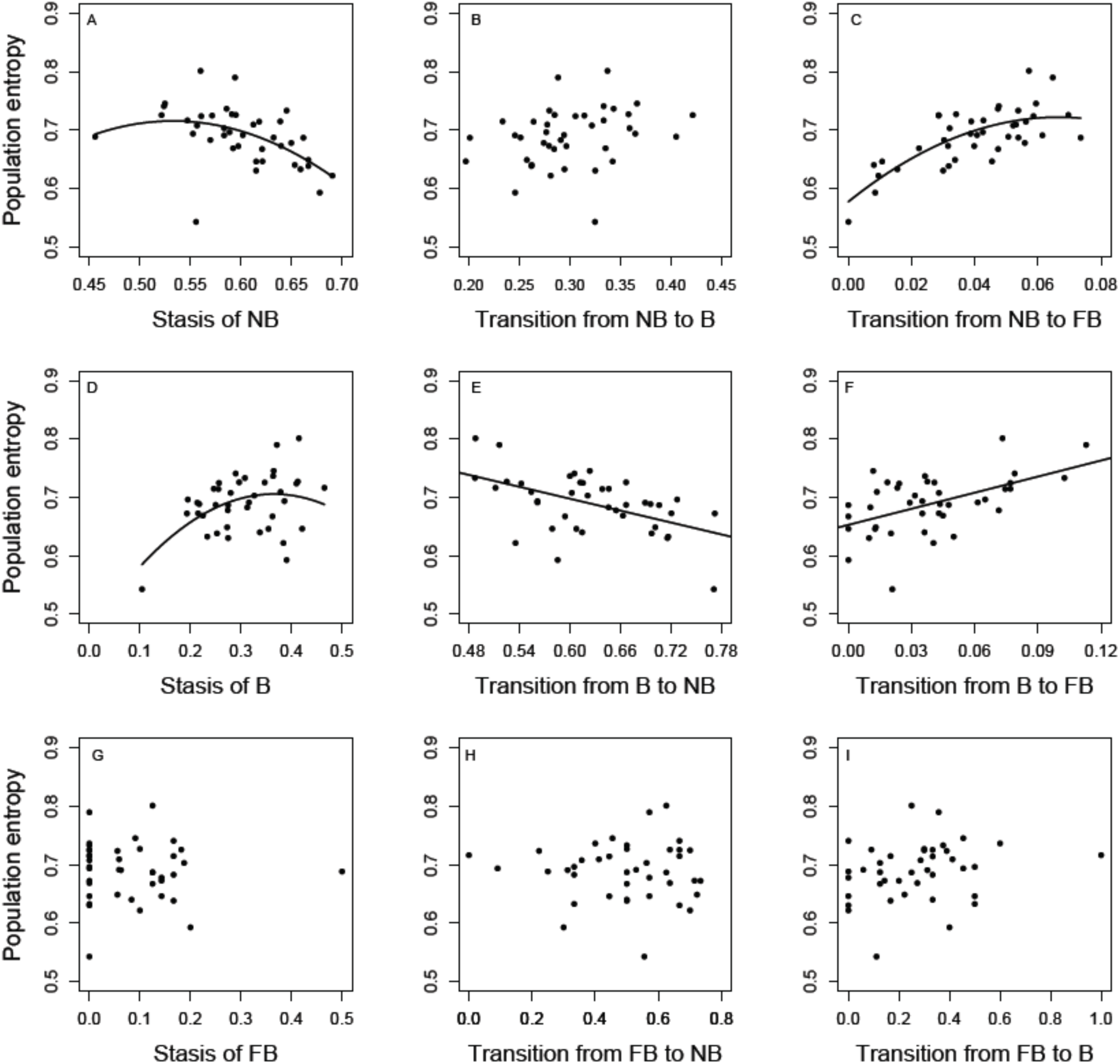
Annual population entropy as a function of stage transitions among mature females.

Our analysis also illustrates that the among year variation in *H*_**R***_t_*_ is mainly driven by stage transitions rather than survival differences among stages (Online Appendix G). Even though we found a strong correlation between the population entropy of matrix **R** and matrix **Pʹ**, only the annual mean survival rates of FB contributed to the variability in *H*_**R***t*_ (Table V). Thereby we indicate that the influence of survival differences among stages on the variability in life courses is negligible (Figure AG 1).

### Stable stage distribution and diversification of life courses

Our annual models mirrored the observed variability in stage distribution across periods, and thus across densities (Figure 3B, 3D). Such observed distribution of females was related to *H*_**R***_t_*_ (Table V). However, *H*_**R***_t_*_ did not relate to changes in density (Figure 4). Thereby we show the independence of the population’s stable stage distribution from its entropy. To further illustrate this independence between the SSD and entropy, we estimated the mean cumulative reproduction (CR) of 100 simulated individual trajectories using two non-absorbing matrices **Pʹ** with similar SSD (1983: NB=0.70, B=0.29, FB=0.01; 1990: NB=0.70, B=0.27, FB=0.03) but different *H*_**R***_t_*_ (*H*=0.60 vs. *H*=0.71, respectively) (Figure 6). Our simulation shows that populations with a similar SSD may still present different amount of individual variation in a specific period in time. Figure 6 shows the population-level effect of such variation in entropy.

**Figure 6:**
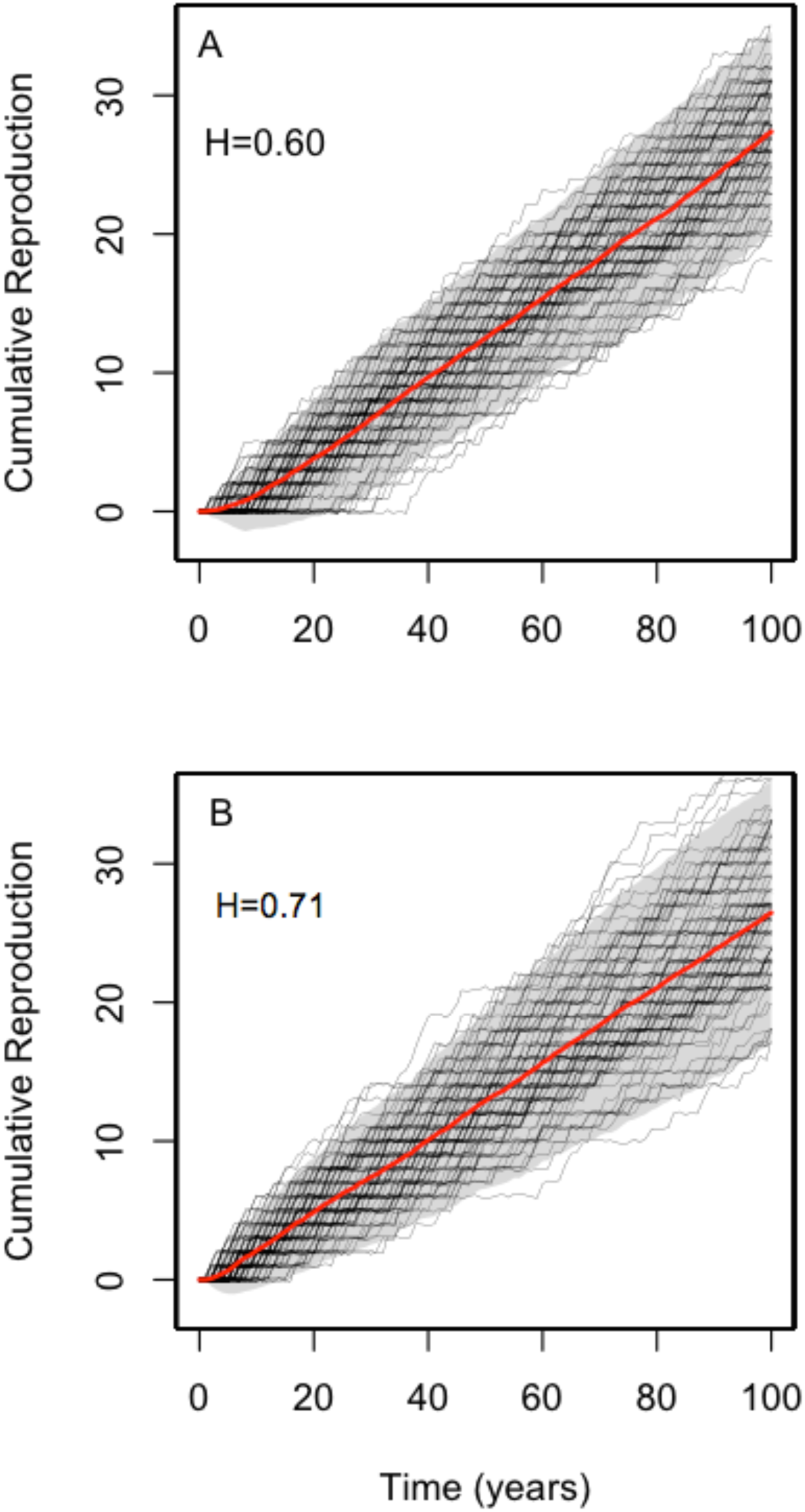
Simulations on cumulative reproduction and age at death for periods of similar stable stage distribution (SSD) and different *H*. (A) represents period 1983 (SSD: NB=0.70, B=0.29, FB=0.01), (B) represents period 1990 (SSD: NB=0.70, B=0.27, FB=0.03). Black line: cumulative reproduction of an individual, red line: mean cumulative reproduction, shaded area: 95% confidence interval.

### Mechanism behind the estimate of population entropy

In order to investigate which stage transition had the highest effect on entropy, and whether or not it relates to density, we performed a perturbation analysis. We found entropy to be highly sensitive to changes between NB stasis and the transition probability of NB to B (Figure 7). Our model with effect of increased NB stasis and reduced probabilities of transitioning to B (scenario ii) was better supported (Figure 7B; Table AH I) compared to the other scenarios where we traded off against FB or a combined decrease of B and FB (Figure 7A, C). Although our perturbation analysis is an oversimplification, it allows us to explore potential changes in *H* given changes in stage transitions.

**Figure 7:**
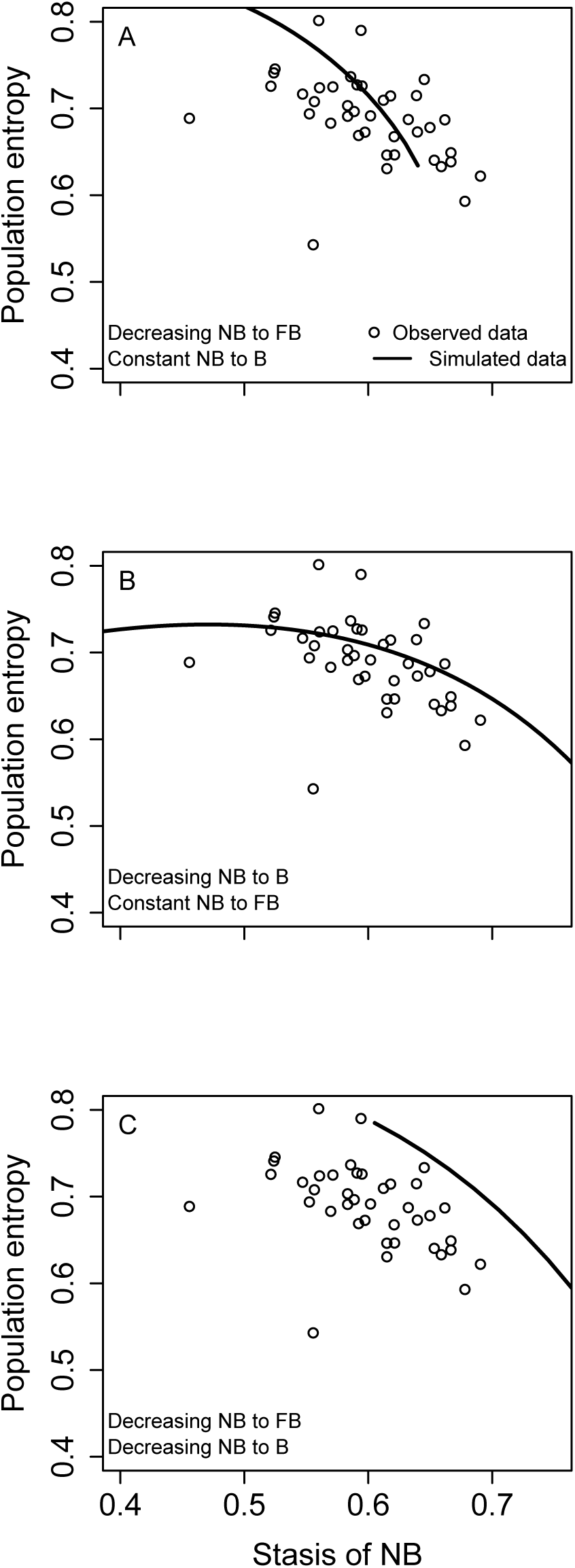
Variability in population entropy generated by simulating an increase in the stasis probability of nonbreeder females given A) a decrease in the transition probability from nonbreeder to failed breeders; B) a decrease in the transition probability from NB to B; C) a proportional decrease in the probability transitions from NB to FB and NB to B. Open circles represent the observed population entropy.

## DISCUSSION

Our analysis on the Cayo Santiago female rhesus macaques reveals a large year-to-year variation in the rate of diversification of individual life courses (population entropy). We demonstrate that such variation is independent of the well known ecological mechanisms underlying structural composition (SSD) in a population; e.g., density-dependence.

Estimating the population entropy, *H*, requires an estimate of SSD in order to weight for the proportion of individuals in each stage. This allows us to account for how many individuals are affected by each transition probability. Entropy relates directly to the probability of an individual to stay in the same stage or move to another, which is independent of the proportion of individuals in such stage (Tuljapurkar et al. 2009). Take for example a 3x3 identity matrix, which has no transitions to other stages, with equal proportion of individuals in each of the three stages (SSD of 1/3 each). Such matrix would have *H* = 0. In contrast, a 3x3 matrix under the same SSD but with equal probabilities in each transition rate would present maximum entropy (Tuljapurkar et al. 2009). This over simplified and non-realistic illustration highlights that SSD and entropy do not have to be directly linked. As a consequence, even if 5% (a very low proportion) of the population is breeding, differences in rates of diversification should be measured by entropy. In the case of the Cayo Santiago macaque population, we found that NB is the most abundant stage among mature female rhesus (>60% in most periods). The annual proportion of NB was affected by density, yet density did not affect *H*; demonstrating the integrative approach that the measurement of entropy encompasses. This holds, as we show, that the amount of variability in life courses depends on all the transition rates, and thus, multiple factors may underlie the estimated individual heterogeneity in a population. In this way, not only our macaque population, but potentially any populations under strong density-dependent regulation, and similar SSD, may still present periods with different rates of diversification in individual life courses. Or in other words, even under strong density regulation, stage dynamics are not fundamentally changed and diverse life histories are not only generated during years of relaxed density regulation.

We found a non-linear relationship between breeders and the total number of adults, indicating that reproduction is boosted as density increases until reaching a maximum from which it is suppressed. The initial boost in reproduction at very low densities might reflect mate-finding limitations (Gascoigne et al. 2009). More likely for Cayo Santiago macaques, low density may boost reproduction at the population level by increasing the probability of females to mature at an earlier age (Bercovitch and Berard 1993), and thus increasing the annual growth rate. In turn, high population density suppressed reproduction. Negative density-dependent dynamics in fertility has been previously described and thought to be driven by social interactions (Hernández-Pacheco et al. 2013). For instance, large group size in primate populations have been related to increased levels of aggression among female primates (Judge and De Waal, 1997), which in turn affect negatively pregnancy outcome (Ha et al. 1999, Ha et al. 2011). The mechanism behind this might include hormonal changes related to stress levels (Ha et al. 1999, Altmann and Alberts 2003), direct aggressive contact inducing abortions, interruption of copulation (Sterck et al. 1997), and harassment by coalitions of relatives (Altmann et al. 1995). Our study suggests such regulatory mechanism, where non-breeders are favored at high densities, might be adaptive under the social stratification of primates. As fewer females in proximity would experience antagonistic encounters among them while mating or during gestation, fewer females would experience hostile environments. Thus, density represents a selective pressure regulating the structural composition of the population (SSD) (and mean vital rates) but does not necessarily regulate the randomness in stage transitions (entropy). Such independence we found provides initial evidence for different evolutionary mechanisms between the ecological density dependent driver and the mechanism that generates the variability in individual life courses selection acts on, but more exploration for detailed understanding is required.

We did not find age specific or temporal changes in stage-specific survival to be related to changes in density nor did they influence *H*. Variability in survival is expected to be low for a provisioned primate population and has been reported previously in Cayo Santiago (Hernández-Pacheco et al. 2013). Cayo Santiago represents an environment of abundant resources given that the amount of food provisioned daily is estimated per capita, no other species is known to compete for resources with the monkeys on the island, and no seasonality in other nutritional resources (e.g., plants, insects) has been reported in the island. Furthermore, populations of social animals may present sophisticated forms of competition that prevent individuals from dying (White 2001). Rather than creating variability in survival, such competition pushes a proportion of individuals (the losers) to remain as nonbreeders (or subordinates), ensuring the effective use of resources. In this way, adult survival among rhesus macaques appears to follow other mammal populations in which selection acts to lower the variance in the population growth rate by minimizing the variation in adult survival (Gaillard and Yoccoz 2003; Hernández-Pacheco et al. 2013). Thus, changes in density do not necessarily translate into survival fluctuations. In populations under such selective pressure, variability in survival is rather minimal and does not represent a significant driver of heterogeneity in individual life courses. The unique characteristics of the population (e.g., food provisioning), coupled to its complex social system, likely account for both the absence of density effects in survival and the weak, but still present, relationship between the population’s structural composition (and reproduction) and density.

Variation in the natal environment can also influence the fate of individuals through early-late life trade-offs (Reid et al. 2003, Plard et al. 2012, Lemaître et al. 2015). For instance, females of a red deer population (*Cervus elaphus*) experiencing high levels of resource competition (high population density) during early life showed faster rates of senescence as adults (lower survival rate) (Nussey et al. 2007). Similarly, female macaques from Cayo Santiago also show a trade-off between age of first reproduction and survival (Blomquist 2009). Here we show that the entropy of a particular cohort is not related to neither the entropy of the year at birth nor to the corresponding density at birth. Our result therefore depicts density at birth as a negligible factor for the diversification of reproductive trajectories later in life.

By us perturbing the stasis of nonbreeders and the transition probability from nonbreeders to breeders we obtained the best-fitted model for the observed annual entropy of the rhesus macaques during the 40-year period. Although this is an oversimplification of the mechanism behind entropy, and our results are subjected to the observed range of densities, the perturbation analysis allows us to explore potential changes in *H* given changes in transitions of the most abundant stage; nonbreeders. Our finding of nonbreeder stasis trading off against breeder probabilities suggests that entropy might be negatively influenced by extreme densities that have not yet been observed in Cayo Santiago. In such scenario, we would expect a reduction in the diversification of life courses (low *H*), being NB the optimal stage.

Our study presents a novel approach to understand how individual heterogeneity is influenced by environmental fluctuations. In particular, it tries to understand if variation in population density influenced individual life history trajectories. Yet, our analysis should be extended to several wild populations with periods of strong density-dependence in order to potentially generalize our conclusions. Our targeted population presents potential limitations for generalization, as wild populations under strong density regulation might experience reduction in entropy with some transition probabilities trading off against others. Yet, Cayo Santiago macaques present an excellent opportunity to address density-effects on the diversity in individual life courses. For instance, management at Cayo Santiago might diminish resource competition at high densities (weak density-dependence). However, food provisioning also buffers fluctuations in the environment that are density-independent, allowing for more accurate detection of density-dependent responses, whether strong or weak (White 2001). Furthermore, as the population is closely managed and thus resources do not vary significantly, we might have not expected the observed density-dependent regulation, nor the substantial variation in entropy. This strongly suggests other forces, rather than density, are acting towards shaping individual variability in Cayo Santiago female macaques, and highlights the appropriateness and unique opportunity this population offers in addressing variability in life histories.

Current research on life history evolution centers on explaining whether the observed life history patterns in a population are adaptive, that is whether patterns result from variation in heritable latent traits fixed at birth, or are rather neutral resulting from stochastic variation in stage dynamics (Tuljapurkar et al. 2009, Steiner et al. 2010, Plard et al. 2012). We have shown here that density-dependent regulation does not necessarily govern the variation in individual life courses on which natural selection can act. Although high population density affected the population structure by reducing fertility, the rate of diversification in individual life courses remained relatively high (*H* > 0.50) across the 40-year period. Thus, our analysis on Cayo Santiago macaques reveals a large variability of unknown source in driving variability in life courses. The causes may lie in differences among the response of individuals to changes in their environment (Plard et al. 2012). Joint models analyzing the contribution of fixed heterogeneity and stochastic stage dynamics (entropy) (Tuljapurkar et al. 2009) across a population density gradient would allow us to test for interactions among genes, neutral variation, and major ecological processes in generating reproductive trajectories among individuals. In this way, we argue that measuring entropy might prove useful in understanding and revealing drivers of diversification in life courses, and helping us to understand whether such diversification is adaptive or neutral.

